# DMD-based super-resolution structured illumination microscopy visualizes live cell dynamics at high speed and low cost

**DOI:** 10.1101/797670

**Authors:** Alice Sandmeyer, Mario Lachetta, Hauke Sandmeyer, Wolfgang Hübner, Thomas Huser, Marcel Müller

## Abstract

Structured illumination microscopy (SIM) is among the most widely used super-resolution fluorescence microscopy techniques for visualizing the dynamics of cellular organelles, such as mitochondria, the endoplasmic reticulum, or the cytoskeleton. In its most wide-spread implementation, SIM relies on the creation of an interference pattern at the diffraction limit using the coherent addition of laser beams created by a diffraction pattern.

Spatial light modulators based on liquid crystal displays allow SIM microscopes to run at image rates of up to hundreds of super-resolved images per second. Digital micromirror devices are another natural choice for creating interference-based SIM patterns, but are not used to their fullest potential because of the blazed grating effect. This effect arises due to the fixed angles between which the mirrors can be switched, creating a sawtooth arrangement of mirrors and thus leading to a change in the intensity distribution of the diffracted beams. This results in SIM patterns with varying modulation contrast which are prone to reconstruction artifacts.

We have carefully studied the blazed grating effect of DMDs by simulations, varying a range of parameters and compared the simulation results with experiments. This allowed us to identify settings which result in very high modulation contrast across all angles and phases required to generate 2-beam SIM pattern. The use of inexpensive industry-grade CMOS cameras as well as low-cost lasers enabled us to construct a cost-effective, high-speed SIM system. Reconstruction of the super-resolved SIM images is achieved on a recently demonstrated parallel-computing platform, which allowed us to visualize living cells with super-resolution at multiple reconstructed frames per second in real time. We demonstrate the versatility of this new platform by imaging cellular organelle dynamics based on live-cell fluorescent stains as well as with fluorescent protein stained samples.

## 1. Introduction

Fluorescence microscopy is a common method to visualize biological and medical samples with molecular specificity. Its spatial resolution is, however, limited due to diffraction, a fundamental physical effect which yields the Abbe law in microscopy, restricting it to about half the wavelength[1].

Therefore, several so-called super-resolution methods have been developed over the last decades to circumvent this limit. Structured illumination microscopy[2, 3, 4] (SIM) is an established technique for super-resolved imaging. Its particular strength is the capability to image at high frame rates and with low phototoxicity, which makes it a highly effective tool for live-cell imaging[5, 6, 7].

Different implementations of SIM exist that can serve as the illumination system for a SIM microscope. Opto-mechanic implementations use a diffraction grating that is either mechanically shifted and rotated[4] or steered by galvanometric mirrors to create interfering light beams and, thus, the SIM pattern in the sample plane. These systems are complex to build, align and maintain, so with the advent of spatial light modulators, these devices were rapidly employed as electronically controlled gratings for SIM[8].

Ferro-electric light modulators (FLCOS devices) are a popular choice to construct especially fast SIM systems [9, 10, 11]. While these devices are successfully used by many groups, they are not without drawbacks. For example, their operating principle does not allow for the continuous display of a pattern, but requires constant switching between a positive and an inverted image. Also, currently only a single supplier offers a limited choice of FLCOS systems.

Digital mirror devices (DMD) are also a promising option for structured illumination [12, 13, 14, 15, 16, 17, 18]. They are available in a variety of models, can provide even faster switching times than the FLCOS system, and can maintain a set pattern for extended durations without the need to switch or refresh the image. Additionally, they are slightly more cost-effective. Their working principle, however, leads to a different distribution of intensities in different diffraction orders when used in conjunction with a coherent laser light source:

The surface of a DMD is formed by a lattice of micro-mechanically driven mirror elements. DMDs are binary in nature, where each element can independently be switched between two states, for example between tilt angles of +12° and −12°, respectively. Typically, these states are denoted as ‘ON’ and ‘OFF’, with those mirrors in the on-position reflecting light in a desired direction, and those in the off-position reflecting towards an absorbing element. In this way, DMDs are seen as amplitude-modulating devices, and for an incoherent illumination source[12], this yields a very effective model of the DMDs operation.

An important aspect comes into play when using a coherent laser light source. Here, the sawtooth-like surface formed by the tilted mirrors leads to a so-called *blazed grating*, which superposes the regular SIM interference pattern generated by the pattern displayed by the device, with the underlying microscopic mirror structure [19, 20]. Thus, for coherent light, the device can be viewed as phase-modulating, because each point on the tilted mirror surfaces introduces a corresponding phase delay.

Only if the blazed grating effect is modeled and properly taken into account, can DMDs be effectively employed for structured illumination microscopy. Here, we explore some of their benefits, by using them to create a rather compact (enabled by the small pixel size) and cost-effective (due to their low price point and overall advances in cost-effective microscopy) SIM system.

Our work aims to provide three results: We modeled, simulated and experimentally verified the blazed grating effect introduced by the DMD in conjunction with SIM patterns. This allows us to employ DMD-devices effectively in coherent SIM applications. We then demonstrate how the advantages of DMD technology can be employed to construct a compact and very cost-effective SIM system. Lastly, we show the performance of such a system by employing it for fixed and live-cell imaging of biological samples.

## 2. Analysis of blazed grating effect: simulation, experiments and results

### 2.1. Coherent illumination of DMDs

An optimal illumination pattern for SIM features high modulation contrast, which directly relates to the order strength in the SIM reconstruction, and thus to the reconstruction quality[21, 22]. As the pattern is generated by interfering two beams of coherent light, these beams have to be of the same intensity and, ideally, the same polarization. Otherwise, the destructive interference is not complete and the modulation contrast is compromised.

This even intensity distribution is difficult to achieve with a DMD. The underlying mirror structure of the DMD results in a tilted reflection grating which leads to the previously mentioned blazed grating effect. Here, the positions of the non-zero diffraction orders relative to the main diffraction order depend on the angle of incidence, grating pitch and wavelength. The envelope of the diffraction pattern and its center also depend on the angle of incidence and tilt angle of the mirrors, whereas the positions of the minima are determined by the effective width of the mirrors and wavelength. Thus, attempts to match one maximum diffraction order, or rather its position, with the envelope center of the overall intensity distribution are complicated by many parameters. In case a match is found, the so-called *blaze condition* is fulfilled and an even intensity distribution of the first diffraction orders is established. If a DMD is used, most of the parameters (mirror dimension, tilt angle, etc.) are readily fixed by the device manufacturer, and therefore only the angle of incidence can be adjusted for any given wavelength to fulfill the blaze condition and to find the so-called *blaze angle* [23].

### 2.2. Effect of underlying blazed grating

First, we investigate the blazed grating effect of the DMD for the case where all mirrors are tilted in one direction and no SIM pattern is displayed by measuring the intensity pattern vs. the angle *α* of the incoming light beam (fig. 1). In order to easily reproduce the positioning of the angle of the incoming light, the laser is coupled into a high-power fiber and then collimated with a custom-made fiber collimator before illuminating the DMD (fig. 1a). Since the mirrors tilt along their diagonal axis, the DMD is also rotated by 45°. Thus, the tilt axis is perpendicular to the plane which is formed by the incoming light and the diffracted light, so that the two-dimensional problem reduces to a one-dimensional problem. Now, the incoming light and the brightest diffraction order form a plane which is parallel to the optical table, so that the subsequent alignment of the microscope system is simplified. A paper screen is used to identify the diffraction pattern and the angle *β* of the maximum diffraction order of the reflected light, which is now referred to as main diffraction order. The intensity of the main diffraction order and the ±1 diffraction orders surrounding it, are measured with a conventional power meter (fig. 1b) to gain some initial insight into the intensity distribution of the envelope function of the diffraction pattern (i.e. its intensity distribution). We evaluated both possible flip directions of the mirrors, the ON (+12°) and OFF (−12°) state and changed *α* from ± 17° to ± 30°, respectively.

**Figure 1:**
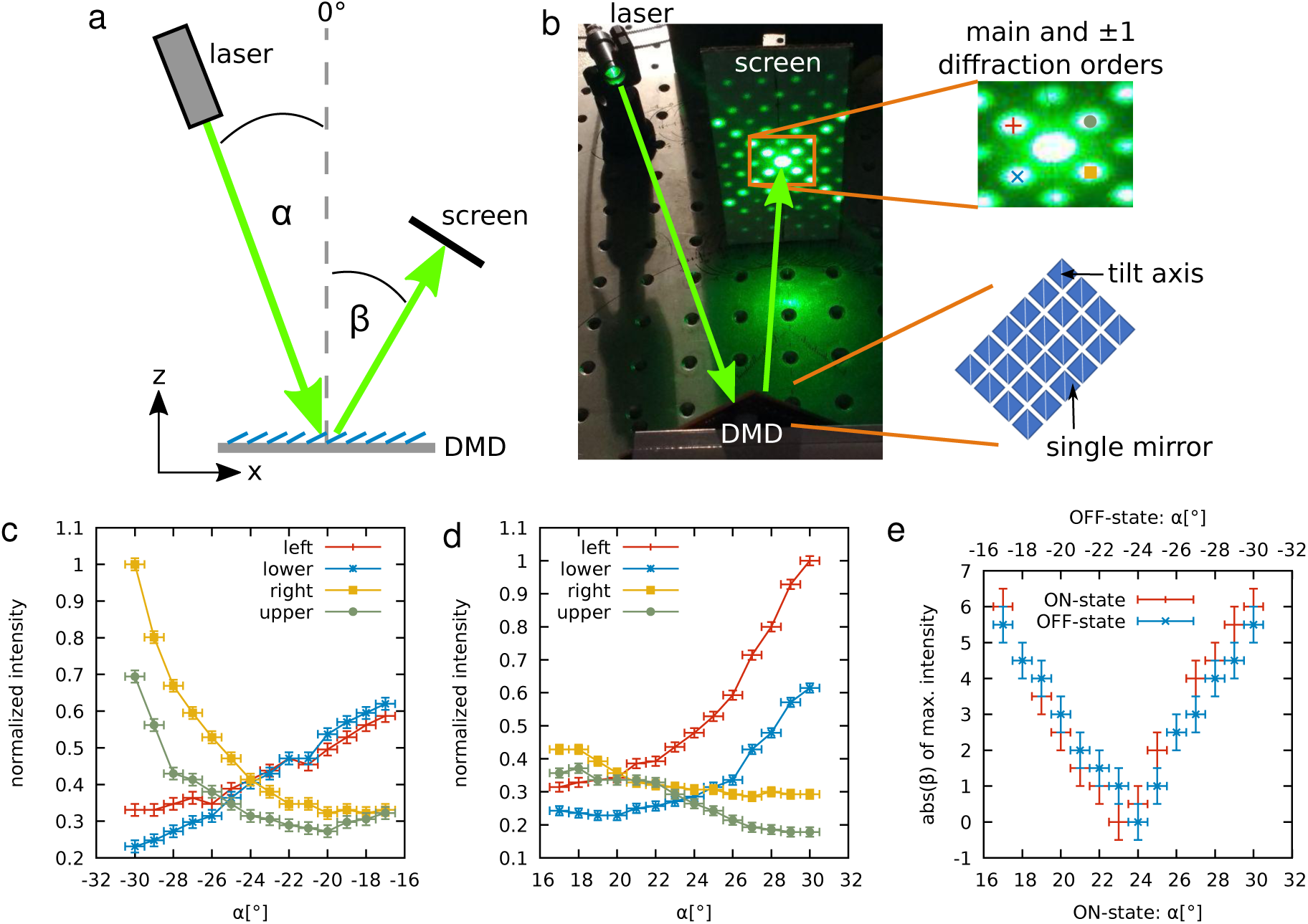
Evaluation of the blazed grating effect underlying a DMD. (a) Schematic drawing of the experiment. All mirrors of the DMD are tilted in one direction, where a tilt angle of +12° represents the ON position and a tilt angle of −12° represents the OFF position. *α* is the incident angle, whereas *β* is the diffraction angle of the main diffraction order. (b) Photograph of the experiment in the laboratory. Here, the entire DMD is rotated by 45°, so that the diffraction pattern is also rotated. The 1 diffraction orders of the underlying structure are labeled with symbols and colors which are also represented in (c) and (d). Normalized intensity measured at the ±1 diffraction orders if all mirrors are switched to the OFF-state (c) or ON-state (d). (e) Absolute value of the diffraction angle *β* of the main diffraction order depending on the angle of incidence *α*.

The measurement of the intensity in the main diffraction order reveals no dramatic difference between the lowest measured value and the highest one, which is about 13 % (ON) and 9 % (OFF) of the full laser power, respectively. Nevertheless, the intensity distribution in the first diffraction orders highly depends on the angle of incidence (fig. 1c and d). If the measured intensity of all first diffraction orders are as close to being equal as possible, we assume that the blaze condition is fulfilled. In case of the OFF-state (fig. 1c) there is an area at roughly *α* = 24.5° ±1° where the intensities are almost equal. However, if all mirrors are flipped to the ON-state (fig. 1d), the result is different from the one obtained for the OFF-state. In addition, it is more difficult to identify the blaze angle because the area where all first diffraction orders have the same intensity is significantly broadened compared to the OFF-state. The absolute value of the diffraction angle of the main diffraction order *β* also reveals no symmetric behavior in the ON- and OFF-states (fig. 1e). Neither the offset nor the slope is equal. Therefore, the tilt angle is possibly not +12° (ON) and −12° (OFF). According to the manufacturer’s data sheet of the DMD, the tilt angle error is precise to within ±1°, and therefore, a symmetric result cannot necessarily be expected.

Nonetheless, the OFF-state appears to be more promising for SIM because based on the results above it will be easier to identify proper values for the blaze angle using this mirror position. This observation does, however, only concern the blazed grating effect of the underlying structure. Further measurements with the SIM pattern on the DMD are more complex if the same range of angles *α* is used. This will allow the use of the results of the underlying structure as start parameters for the search of the blaze angle. To obtain the exact values and then confirm these results, more calculations and simulations are needed as detailed below.

### 2.3. Simulation of the blazed grating effect

To better understand the results obtained by measurements, we chose to model the DMD and simulated the diffraction pattern depending on the angle of incidence *α* [24]. To accomplish this, the DMD needs to be described mathematically, and the electric field reflected from its surface has to be calculated for different positions and states of the mirrors. Finally, the blaze angle has to be determined for the case, where a SIM grating pattern is displayed on the DMD.

Since the DMD is basically a two dimensional array of single mirrors, we first modeled a single mirror using a suitable coordinate system (fig. 2a). We assumed that each non-tilted mirror is in the xy-plane of a Cartesian coordinate system. To describe a point on a single mirror 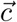, we introduce

**Figure 2:**
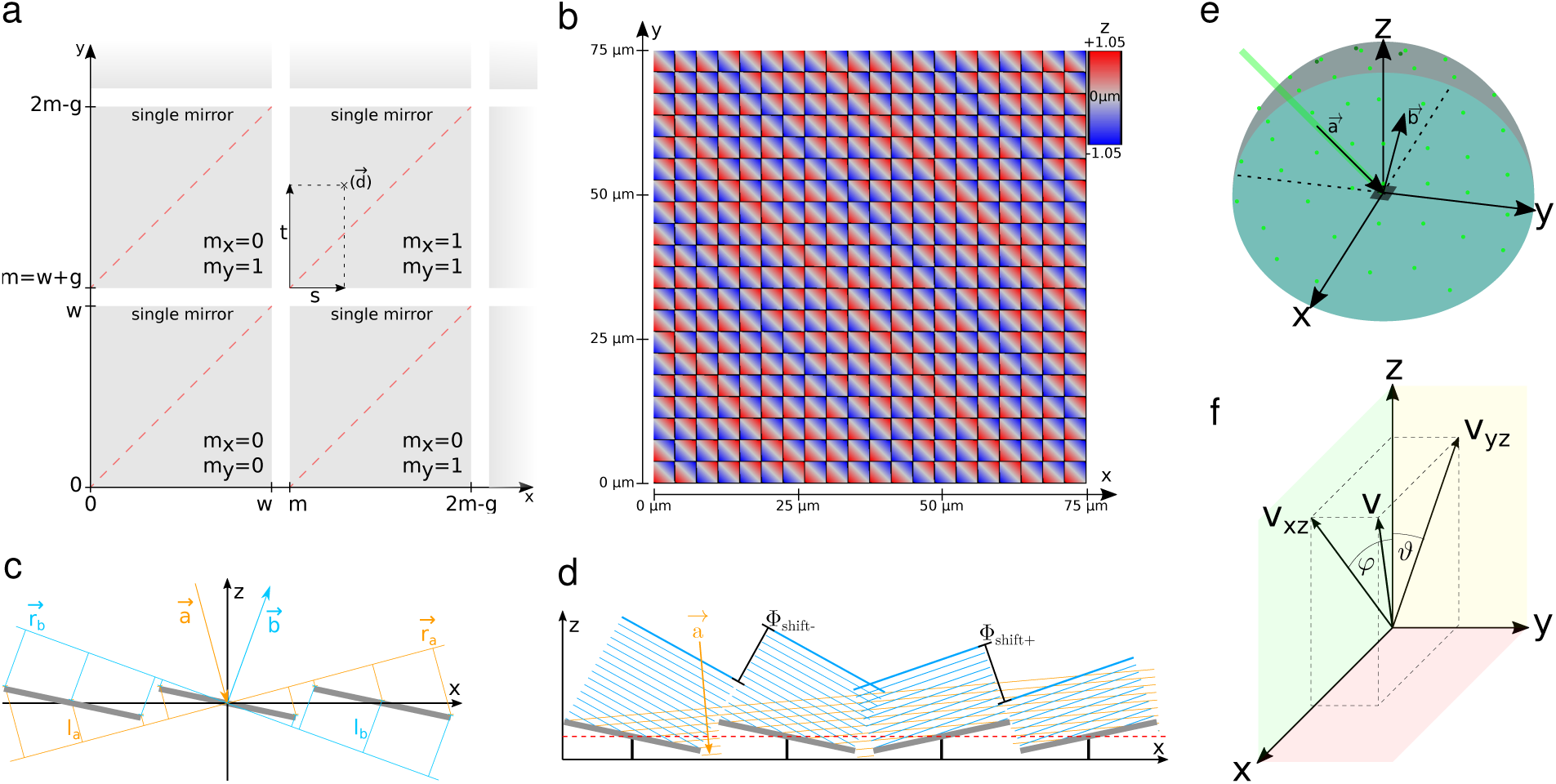
Modeling (coherent) light propagation on a DMD surface. (a) Physical dimension and rotation axis of the micro-mirrors. (b) Resulting height profile of the DMD surface, which corresponds to phase delay. (c) Phase shift of different points in a planar wave-front, propagating towards the DMD in direction 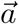, or reflected from the DMD in direction 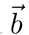, see eqs. (7) and (8). (d) Simplified modelling of light wave-fronts being reflected by the micro-mirrors, where a single, analytically known wave-front emanates from each mirror. This approach is correct in far-field / Frauenhofer approximation. (e) Visualization of light input and output vectors in the 3-dimensional space of the DMD. (f) Visualization of the tip/tilt angular coordinate system used to represent directionality of e.g. the directions 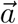 and 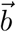.

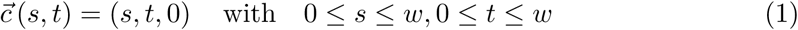

where *w* is the width of one mirror, *s* and *t* are the points in x- and y-direction, respectively. Now, we need to rotate the mirror along its diagonal axis 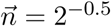 with the rotation matrix

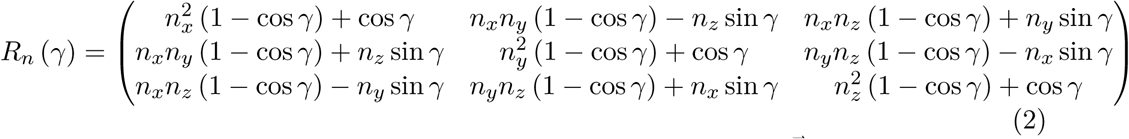

where *γ* is the tilt angle along the diagonal. To describe a point 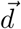 on the surface of the DMD, each mirror has to be addressed by introducing the grid 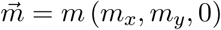,where *m*_*x*_ ∈ℕ_0_ and *m*_*y*_ ∈ℕ_0_, and the gap *g* between the mirrors is considered in *m* = *w* + *g*. With eq. (1) and eq. (2), the surface is now parameterized by rotating a single mirror and placing it to its position

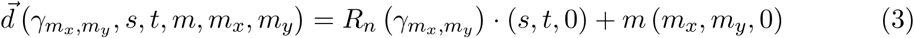

Here, *γ* is now expressed as 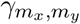 to address each mirror. The resulting surface of the DMD can then be modulated and visualized with the tilt of the mirrors (fig. 2b).

Next, we need to calculate the diffracted intensity distribution *I* which is the square of the absolute value of the reflected electric field *E*. Since we use monochromatic and coherent light, the time-independent electric field can be described as

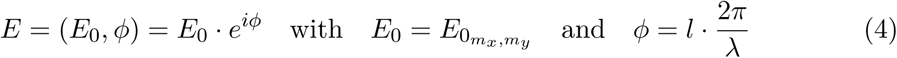

where *E*_0_ is the amplitude and *ϕ* the phase shift. The Gaussian beam profile of the laser beam is approximately constant for a single mirror, and therefore *E*_0_ depending on *m*_*x*_ and *m*_*y*_ needs to be introduced. The phase shift *ϕ* is determined by the path length *l* and the wavelength *λ*.

However, since we use the case of Fraunhofer diffraction for this simulation, the diffracted light spreads out in all three spatial dimensions in a spherical way (fig. 2e), so that it is more convenient to change the Cartesian coordinate system to one where the unit vector 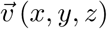 is now associated with 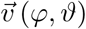 (fig. 2f). The propagation vectors of the incoming light 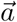 and the diffracted light 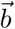 are normalized and can be described as follows

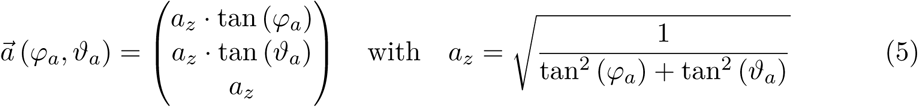

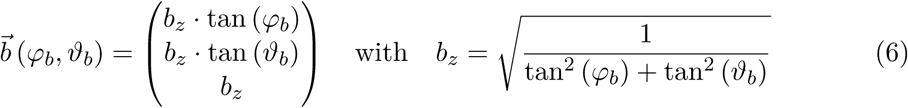

The propagation vector of the electric field illuminating the DMD 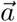 is perpendicular to the wavefront plane 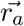, which is also true for the propagation vector of the diffracted electric field 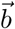 and its wavefront plane 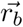 (fig. 2c). Now, we set the wavefront plane in the origin of the coordinate system, so that the Hesse normal form of the wavefront is

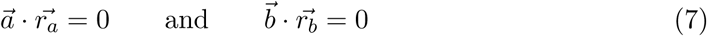

Due to the tilted mirrors, the planar wavefront, propagating along the direction 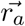 hits each point of the DMD with a different phase shift, which results in a different path length *l*_*a*_. By calculating the distance of the fixed wavefront plane 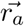 eq. (7) to the DMD 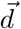,we obtain *l*_*a*_, which can be seen as a distance from a point to a plane. Additionally, the diffracted light also contributes an extra path length *l*_*b*_ (fig. 2c). Due to the mathematical determination of distance point-plane, *l*_*a*_ and *l*_*b*_ can be expressed as

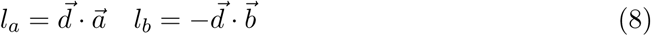

Thus, the total path length *l* is the sum of *l*_*a*_ and *l*_*b*_, which we can insert in eq. (4), and therefore we get a phase shift of

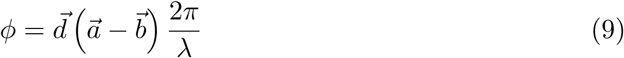

and an electric field of

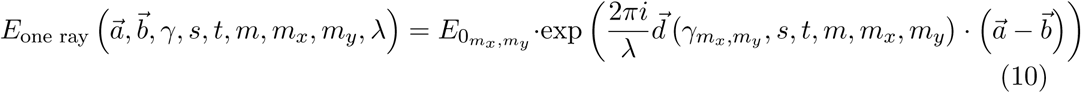

At this point, we can see why in this idealized model, the DMD acts almost purely as a phase modulator. For all wavefront components hitting a mirror, only the phase is affected by the tilt of these mirrors. The polarization of the light is not modified, while the amplitude is modulated in a binary fashion, with some components hitting the mirrors being unmodified, and some components missing a mirror element being absorbed entirely. Now, we need to consider each point of a single mirror, so that an integration of *s* and *t* is necessary. Additionally, each mirror of the DMD contributes to the resulting electric field, so that a sum over each mirror is required. Thus, by modifying eq. (10) we get

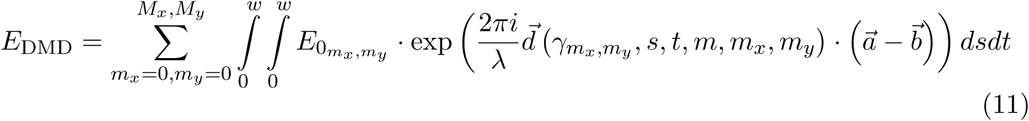

where *M*_*x*_ and *M*_*y*_ are the total numbers of mirrors in x- and y-direction, respectively.

However, we can use several assumptions to solve eq. (11). First, as already mentioned, 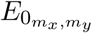 is constant for a single mirror, and therefore it can be extracted from the integral *F*, which we define as

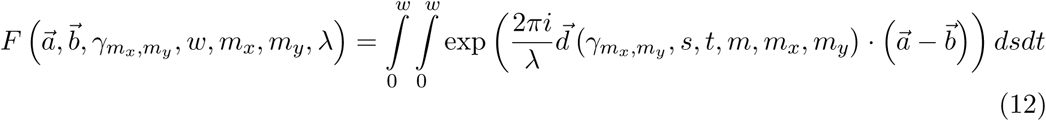

This integral can be solved analytically e.g. by using a computer algebra system (see source code repository accompaning the manuscirpt). Nevertheless, eq. (12) can also be simplified to obtain more insight into the phase modulation effect. Each mirror tilts by either −*γ* or +*γ*, which results in only two different phase shifts, namely *ϕ*_*shift*_ and *ϕ*_*shift*+_ measured relative to a reference mirror (fig. 2d). Thus, we only need to determine two integrals 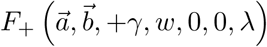 and 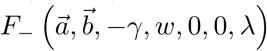 in the origin of our coordinate system and multiply a complex factor due to the mirror position which can be seen by inserting eq. (3) in eq. (12):

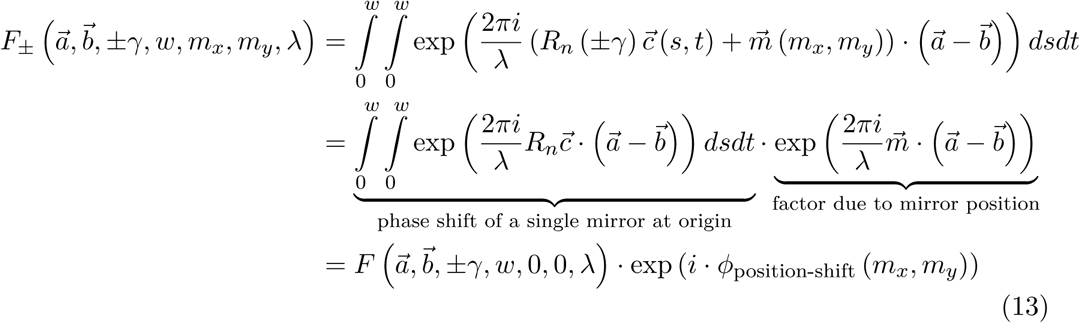

Finally, we can calculate the intensity distribution by inserting eq. (13) in eq. (11) and determining the square of the absolute value

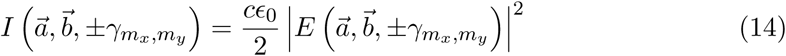

with *c* the speed of light and *ϵ*_0_ the vacuum permittivity.

However, eq. (14) gives us several possibilities for different experimental realizations. 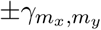 allows us to modulate the DMD in a large number of different configurations. We can define the number of mirrors, the tilt angle and even display the SIM pattern, because each mirror is addressed individually. The incident angle, which is crucial for an even intensity pattern, can be chosen by setting 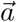 properly. Then, we need to sample the space by varying 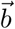 to determine the intensity *I* at each spatial point. We created a software package to numerically perform these calculations, see appendix B.

In our first implementation of the simulation we modeled a single mirror which is either tilted in the ON- or OFF-state and set an angle of incidence of 0° (fig. 3a). The respective intensity patterns display the expected mirrored distribution. In both cases the main diffraction maximum is clearly visible and the first diffraction orders are lying in the x- and y-direction. Therefore, only one state is further investigated.

**Figure 3:**
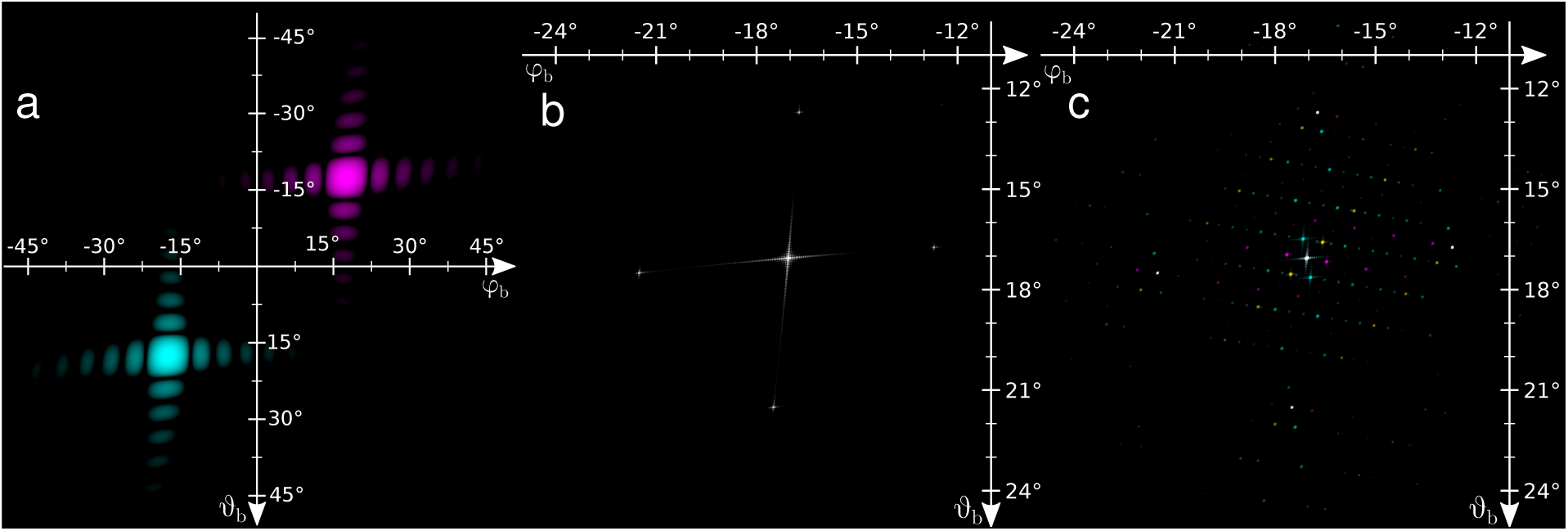
Wave propagation and interference pattern simulation of a DMD surface. A fully coherent light source is simulated illuminating the DMD head-on (angle of in-cidence 0°) and with a Gaussian intensity profile (b and c). The diagrams show the reflected intensity plotted against the output angle (fig. 2), on a logarithmic scale. (a) Diffraction pattern of a (hypothetical) single mirror, in its two possible tilt states (“teal” is OFF and “magenta” is ON). (b) Diffraction pattern of the DMD surface when all micro-mirrors are oriented into the same OFF position. (c) DMD displaying a typical SIM pattern, while illuminated with coherent light. The full diffraction pattern arises from the inert diffraction properties of the DMD (mechanical mirror size and tilt) with the pixelated and binary SIM pattern superimposed. The underlying structure of the mirrors is the OFF position. The three colors indicate a different SIM pattern for each typical angle used for SIM.

In the next iteration of the simulation we modeled a two-dimensional array of mirrors. Here, we chose an array of 100 ×100 mirrors which are all positioned in the OFF-state (fig. 3b). Since the main diffraction order is much brighter than the first diffraction orders, we chose a logarithmic scale to display the data and, therefore, the maxima look like crosses rather than round spots (which is the typical result for linearly scaled axes). Nevertheless, a comparison to the experimental results indicates that these simulated results are, indeed, valid.

Next, we modeled the case where a typical SIM grating pattern is displayed on the simulated DMD with the OFF-state as underlying structure (fig. 3c). In addition, we rotated the SIM grating pattern by two further angles (approx. 60°), so that we obtained the typical three angles that are used for space-filling 2D and 3D SIM. The result, as shown in fig. 3c, indicates that the diffraction orders of the underlying structure are, indeed, modulated and we obtain further diffraction maxima. Also, as can be seen in fig. 3c, the diffraction orders of the underlying structure stay at their original position, although a different pattern is being displayed on the DMD.

So far, we have demonstrated that the SIM pattern in the Fourier plane can be simulated if a DMD is used as the primary device to create interference patterns and we have verified the simulation results with experimental measurements. Next, all possible illumination conditions and their resulting blaze angles need to be identified in order to guide the experimental implementation. Therefore, we used our model to vary between a range of angles of incidence *φ*_*a*_ and *ϑ*_*a*_ and to calculate the resulting intensity patterns. To determine the best candidates for the blaze angle, a comparison of the intensity of the first diffraction orders was chosen as this parameter indicated that it produces sufficiently close results (fig. 4a-c). If the intensities of the first diffraction orders are equal, it indicates that the blaze angle was found. This parameter is, however, specific for one SIM pattern angle but will likely be different for other pattern angles as can be seen in fig. 3c. This behavior can be contributed to the fact that the diffracted intensity pattern of a single mirror is the envelope function of the overall diffraction pattern (fig. 4a). Therefore, it is necessary to match the maximum of the envelope function with the maximum of the main diffraction orders of the SIM pattern in order to obtain an even intensity distribution in all first diffraction orders that are used to create SIM patterns for all SIM pattern angles. To match the results we calculated the distance between the maximum values. Since this process needs to be repeated for thousands of data sets running through all possible variations of angles, we implemented the simulation on a graphics card to accelerate the calculations. The result is shown in fig. 4d and it implies that there are a large number of potential blaze angles according to the dark rings in the figure, which indicate a good match.

**Figure 4:**
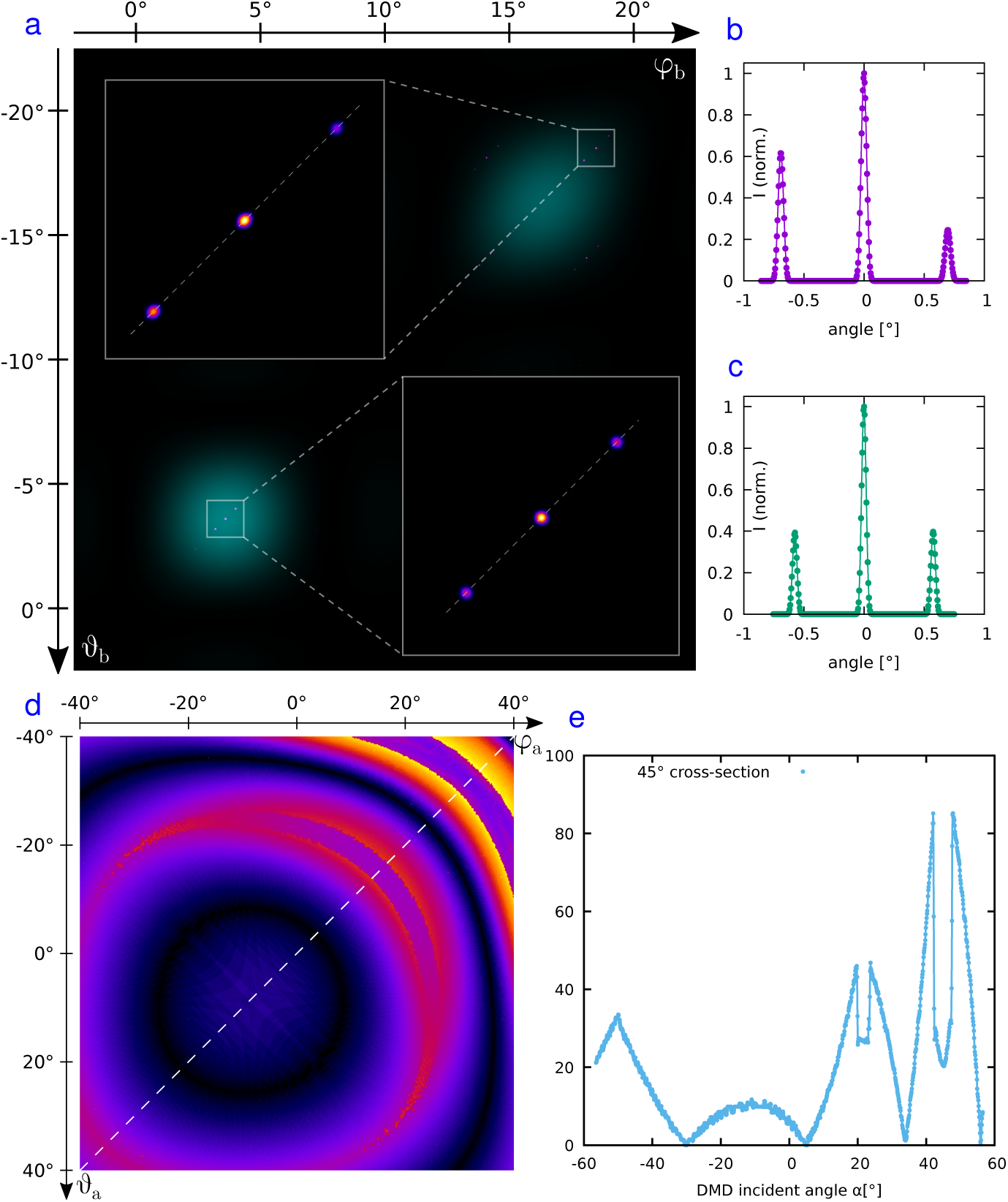
Simulations of DMD wave propagation for optimized use with SIM. For all panels the OFF-state is the underlying structure. (a) Alignments of the SIM pattern (magenta/blue/white) to the diffraction envelope (teal) caused by a single mirror in the OFF-state (fig. 3a). In the top row, the main diffraction order of the SIM pattern does not coalign with the envelope, with yields an uneven intensity distribution in the ± 1 side orders that create the SIM pattern (b). In the bottom row, the main SIM diffraction order coaligns with the envelop, with yields an even intensity distribution into the ± 1 side orders (c). Axis are centered on the main SIM diffraction order and scaled in small angle approximation (b and c). Misalignment between main SIM diffraction order and maximum of the envelop, depending on the input angles (d) and along a 45° cross-section (dotted line, plot in panel e). Alignments close to zero yield equal intensities in the SIM side orders. The 45°line was chosen as it allows for an easy alignment of all angles in one plane (DMD mounted to a table).

To better correlate these results with experiments, the entire simulated DMD structure was rotated by 45°, such that now only the cross-line, where *φ*_a_ = −*ϑ*_a_, is of interest. Additionally, the results shown in fig. 1c and fig. 1d imply the use of the OFF-state as the underlying structure, and therefore this is also used for the simulations. Angle calculations due to different coordinate systems are needed to allow for a proper comparison to the experimental results. At the 45° cross-section, *α* is determined by

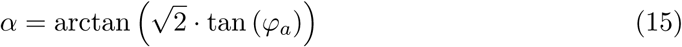

From fig. 4e it is apparent that minima at *y* = 0 occur at several points which indicate the spots where the blaze condition is fulfilled. For the DMD used in our experiments, we found that the blaze angle is at *α* = 27.9° (fig. 4e). As a control, we took a line plot across the main and first diffraction orders to underline our result (fig. 4c). Also, in fig. 4a we can see that the center of the overall mirror envelope is an excellent match with the main diffraction order. To demonstrate a non-matching case, we chose an incident angle of *α* = 53° and the difference in the first diffraction orders is clearly visible from fig. 4b.

### 2.4. Comparison of simulations and experimental results

The predictions made by our simulated DMD data where an actual SIM pattern was modeled have to be verified with experimental data. To facilitate this, we projected the experimentally obtained intensity pattern diffracted by the DMD onto a camera chip using a single lens. This arrangement guarantees that the Fraunhofer diffraction condition is also fulfilled in the experimental case. The camera is positioned in the Fourier plane of the setup. In addition, the power of the laser beam used to illuminate the DMD has to be reduced by neutral density filters to avoid saturating the camera pixels. For the measurements all nine SIM patterns (three illumination angle and three associated phase shifts) were displayed by the DMD with a display time of 105 *µ*s for each single frame. The exposure time of the camera was set to 5 ms. To find the blaze angle the incident angle *α* was set to −24° for the OFF case and to +25° for the ON case according to the results of fig. 1c and fig. 1d. To cover a wide range of angles *α* the DMD was gently tilted by ±0.5° and ±0.3° along the x- and y-direction between measurements using micrometer screws, respectively. This also allowed us to evaluate how critical the precise alignment of the blaze angle is in order to obtain the highest possible SIM modulation contrast.

We started by comparing the distribution of the diffraction orders in fig. 5a (ON) and b (OFF). As expected, the distribution in both cases is very similar, but it looks different when compared to the results obtained by the simulation (fig. 3c and fig. 6f).

**Figure 5:**
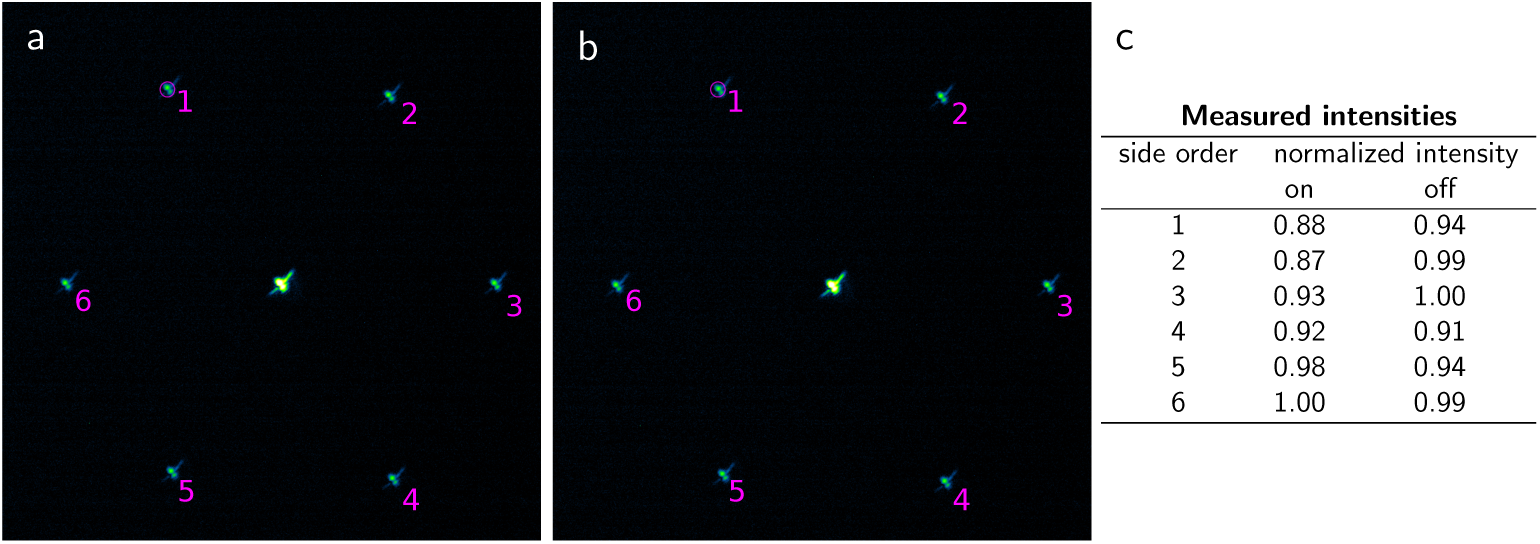
Measured diffraction intensity pattern in the Fourier plane with the SIM pattern displayed on the DMD. With a lens the intensity pattern is focused on a camera chip. In both cases (a-ON and b-OFF, exposure time 5 ms, DMD display time 105 *µ*s) the measured blaze angle is chosen as the incident angle, which is +25.3° for ON and −23.9° for OFF. The difference compared to the simulated data is due to a different tilt angle *γ*. For a better visualization a log scale was set. The mean value of the circled area in (a,b) was analyzed to evaluate the quality of the found blazed angle (c).

**Figure 6:**
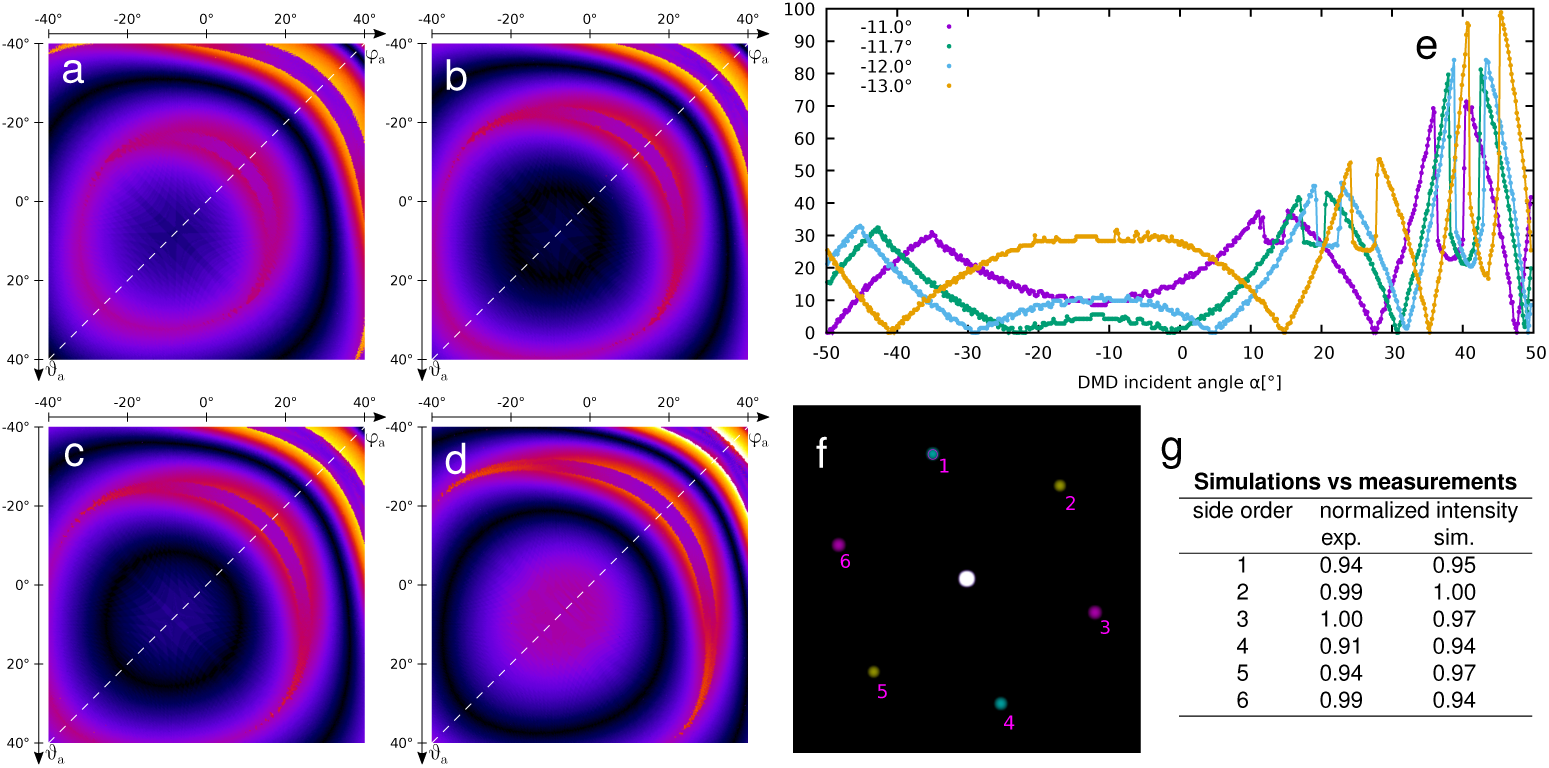
Evaluation of the dependence of the blaze angle on the tilt angle *γ* of the DMD mirrors. Misalignment between main SIM diffraction order and maximum of envelope depending on *γ* (a: − 11°, b: −11.7°, c: −12°, d: −13°). In each simulated figure the incident angle has been changed in order to find the blaze angle. Again, a crosssection at 45° was chosen to compare the simulations results (e) to the experimental data. The blaze condition at *y* = 0, where the plotlines come down to the x-axis, shifts clearly due to small changes of *γ*. For a direct comparison to the experimental results (*α* = −23.9°, *γ* = −11.7°), the same case is simulated (f). Again, a log scale was set for a better visualization. Same colors of the first orders belong to the same SIM angle. The mean of the side orders in the displayed ROI is listed and directly compared to the experimental data of fig. 5c (g).

The characteristic cross shape from the simulations is not visible, instead a diagonal line can be seen. We attribute this to two factors: first, the physical DMD microstructure might deviated from the ideal structure that was used in the simulations. Also, the DMD chip is covered by a protective glass plate, which we did not take into account in the simulations. Indeed, if a different spot of the DMD is illuminated, the diagonal line also moves. This leads us to assume that the glass cover causes a change in the light path and accordingly changes the distriution of the diffraction orders. Additional analyses, such as varying the beam diameter and inserting a Fourier mask, showed that the spots then change to a spherical shape. This is an acceptable solution since in most SIM setups a Fourier mask is used to reject other diffraction orders, anyways.

The blaze angles for our DMD were experimentally found to be *α* = +25.3° in the ON case and *α* = −23.9° in the OFF case (fig. 5a and b). Since a high modulation depth is desired for best SIM performance, the intensity of the first diffraction orders belonging to the same SIM angle is supposed to be equal. We measured the mean intensity values of the first diffraction orders by choosing a region of interest (ROI) across each spot and evaluated their brightness (fig. 5c). For a fair comparison, only the intensities of the same SIM angles need to be compared, namely 1 and 4, 2 and 5, 3 and 6. We found the maximum difference to be 11 % (ON) and 5 % (OFF) for these spots - which remains very similar if the DMD is tilted in x- and y-direction by ±0.5° and ±0.3°. Nevertheless, the absolute values of the experimentally found blaze angels are not the same as those obtained by the simulation. We have to keep in mind, though, that the simulated results were obtained using a fixed tilt angle of *γ* = − 12°. According to the manufacturer, however, *γ* varies from −11° to −13° in the OFF case. Therefore, the impact of different *γ* values on the blaze angle needs to also be investigated.

In order to analyze the dependence of the blaze angle on the tilt angle *γ* of the underlying structure, various cases of *γ* were chosen for the simulations. Again, the distance of the envelope of one diffracting mirror to the main diffraction order (as shown in fig. 4d) has been determined for different angles of incidence to evaluate the blaze angle. As can be seen from fig. 6a to fig. 6d, a small change of the tilt angle leads to rather significant changes in the graphs, resulting in very significant shifts of the blaze angle. Again, only the OFF case as underlying structure is simulated and shown due to the fact that the ON case should be a mirror image of this case. The dark rings which indicate solutions for the blaze condition have very different diameters. As we saw previously in fig. 3d with *γ* = −12°, there are two solutions rings. If the tilt angle decreases to *γ* = −11°, the diameter of the two dark circles also decreases and the inner circle even disappears entirely (fig. 6a). An increase of *γ* to −13° leads to large diameters of the dark rings (fig. 6c). Since the *γ*-range extends from −11 to −13°, and the position of the blaze condition shifts accordingly, a potential maximum value for the blaze angle can be determined. To find this value, the cross-section along the 45° axis, where *ϕ*_*a*_ = − *ϑ*_*a*_, was plotted for the different cases of *γ* (fig. 6e). According to the simulation results a fairly wide range of values for the blaze angle is covered. For example, if *γ* = − 13°, one possible corresponding blaze angle is *α* = −41°.

Based on our simulations we found that a tilt angle of *γ* = −11.7° for the underlying structure, leads to a blaze angle of *α* = −23.9° for the OFF case (fig. 6e). This matches perfectly with the experimentally measured blaze angle and the possible *γ*-range according to the manufacturer. Also, the experimentally measured blaze angle for the ON case, *α* = +25.3°, is consistent with the simulations, since a potentially wide range of blaze angles is covered due to the range of possible *γ* values. To further support our results, we show the simulated diffraction pattern if the tilt angle *γ* were −11.7° and the angle of incidence *α* = −23.9° is the same as the experimentally measured blaze angle for the OFF case (fig. 6f). Again, as in fig. 5, we basically chose the same ROIs covering the first diffraction orders and evaluated the mean values. A direct comparison shows that the simulated results are consistent with the experimental results (fig. 6g).

## 3. Construction of a compact and cost-effective SIM system

Now that we have determined the correct blaze angle, constructing the remaining components of the DMD-SIM microscope is straight-forward. Compared to other SLM-based SIM microscopes, using a DMD is more cost-efficient and allows for a compact design due to the small pixel size of the DMD mirrors. This allowed us to construct a SIM microscope with a rather small footprint, which could be placed on a 40 cm x 90 cm optical bread board (fig. 7a). In addition, we exchanged the laser and the camera from scientific to industry-grade components (see appendix A.2). The laser is typically used for light show applications and the industry-grade CMOS camera has already been tested for other implementations of super-resolution microscopy, where its use has lead to sufficient results as long as the sample brightness does not require single photon counting [25, 26]. Overall, the use of these components allowed us to keep the total cost of the system below 20 k€which is more than ten times less expensive than commercial solutions.

**Figure 7:**
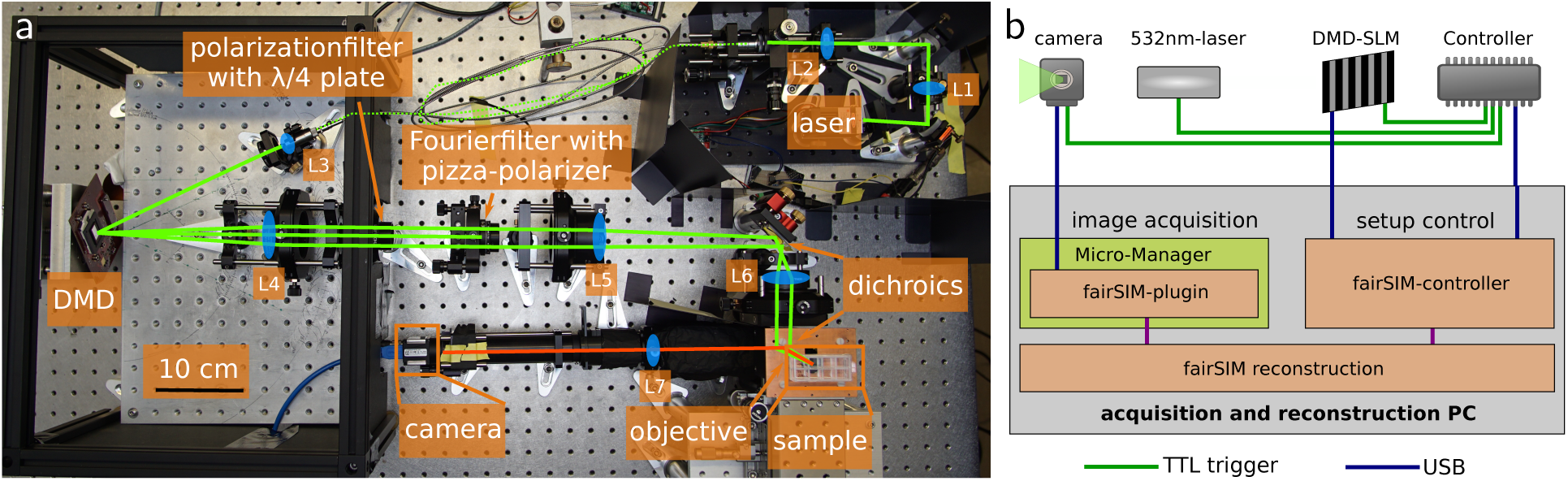
Photograph of the custom-made SIM microscope and schematics of its data stream. (a) A 532 nm laser beam (green) is coupled into a fiber before illuminating the DMD at the blaze angle. Polarization optics are required to achieve the same linear polarization in the corresponding SIM angles. Two dichroic mirrors from the same batch are used to maintain the polarization state and to separate the excitation light from the fluorescence (red) signal. The same objective lens is used to excite the sample and to collect the fluorescence signal which is detected with an industry-grade CMOS camera. (b) Timing control and image data flow. Binary SIM pattern are stored on-board the DMD and selected in sequence by sending TTL trigger pulses to the DMD control board. The DMD, camera and laser light source are synchronized using a micro-controller. The raw image data is fed into a GPU-based real-time data processing system [11]. A detailed description of (a) and (b) can be found in appendix A.1.

The laser is coupled into a high-power single-mode fiber to clean up its beam profile and to enable its easier adjustment of the illumination blaze angle with respect to the DMD surface. The grating patterns displayed on the DMD are calculated via a (modified) search algorithm used to find suitable binary gratings[9, 11, 27]. The light diffracted off of the DMD is subsequently collected by a lens and the polarization of the excitation beam is set by a segmented polarization filter (fig. 7a). The six beams used for generating the SIM interference patterns are then filtered with a custom-made Fourier mask (pinhole pattern). 8 % of the laser power incident on the DMD are ultimately maintained in the two corresponding SIM beams. The same objective lens is used to generate the SIM pattern at the sample location and to collect the fluorescence signal. Since the 532 nm laser can excite a number of *orange* and *red* fluorescent dyes, two detection filter sets are used according to the fluorescence emission. Lastly, the industry-grade camera is used to detect the fluorescence signal with tube lens L7 which results in a projected pixel size of 70 nm and 35 nm for SIM (more details in appendix A.1 and appendix A.2).

To obtain the best possible modulation depth for SIM illumination the relative polarization of the excitation beams is a critical factor. Ideally, the two corresponding beams should have the same linear polarization [28, 29]. Therefore, several components to control the polarization of the excitation beams are utilized in the setup (fig. 7a). First, the diffracted light is elliptically polarized and a linear polarizer in combination with a *λ*/4 plate is used to change the polarization to circular polarization. Right behind the Fourier mask, a segmented pizza-polarizer is placed in the Fourier plane to ensure proper linear polarization of the corresponding beams. Using a perpendicular arrangement of two dichroic mirrors from the same batch, the polarization adjustment is maintained[10].

Acquiring SIM images with high frame rates requires precise timing between the DMD, the laser light source and the camera. The DMD control board allows to upload predefined sequences of pattern, which are displayed with either fixed timing or in reaction to external trigger inputs. The camera equally allows for external triggering of the start of an image exposure, and the laser light source can be switched at high speeds. An inexpensive micro-controller is used as a master clock device to generate these timing pulses.

The raw images acquired by the camera are fed into a real-time processing pipeline (called fairSIM-VIGOR) [11]. Compared to its original implementation, this microscope only needs to process a single color channel with reduced frame rate, so a single computer equipped with a graphics processing unit (GPU) suffices to handle data acquisition, processing and storage. Due to the real-time processing of the image data the user obtains immediate feedback in super-resolved images and can directly evaluate the usability of the data during the experiment.

To test the functionality of the DMD-SIM setup, we imaged TetraSpeck beads (TS) with a diameter of 200 nm and then reconstructed the frame set with fairSIM [30, 11]. For the reconstruction we used a calculated optical transfer function (OTF) and OTF attenuation to suppress out-of-focus fluorescence signals[33, 34]. Besides the summed up wide-field image (WF), we also generated a filtered wide field image (fWF) by applying the generalized Wiener filter step inherent to SIM reconstructions also to the WF data. This is important for a fair visual comparison, as the fWF image provides better contrast, but only the SIM image can truly reveal more details and separate individual beads (fig. 8a-c).

**Figure 8:**
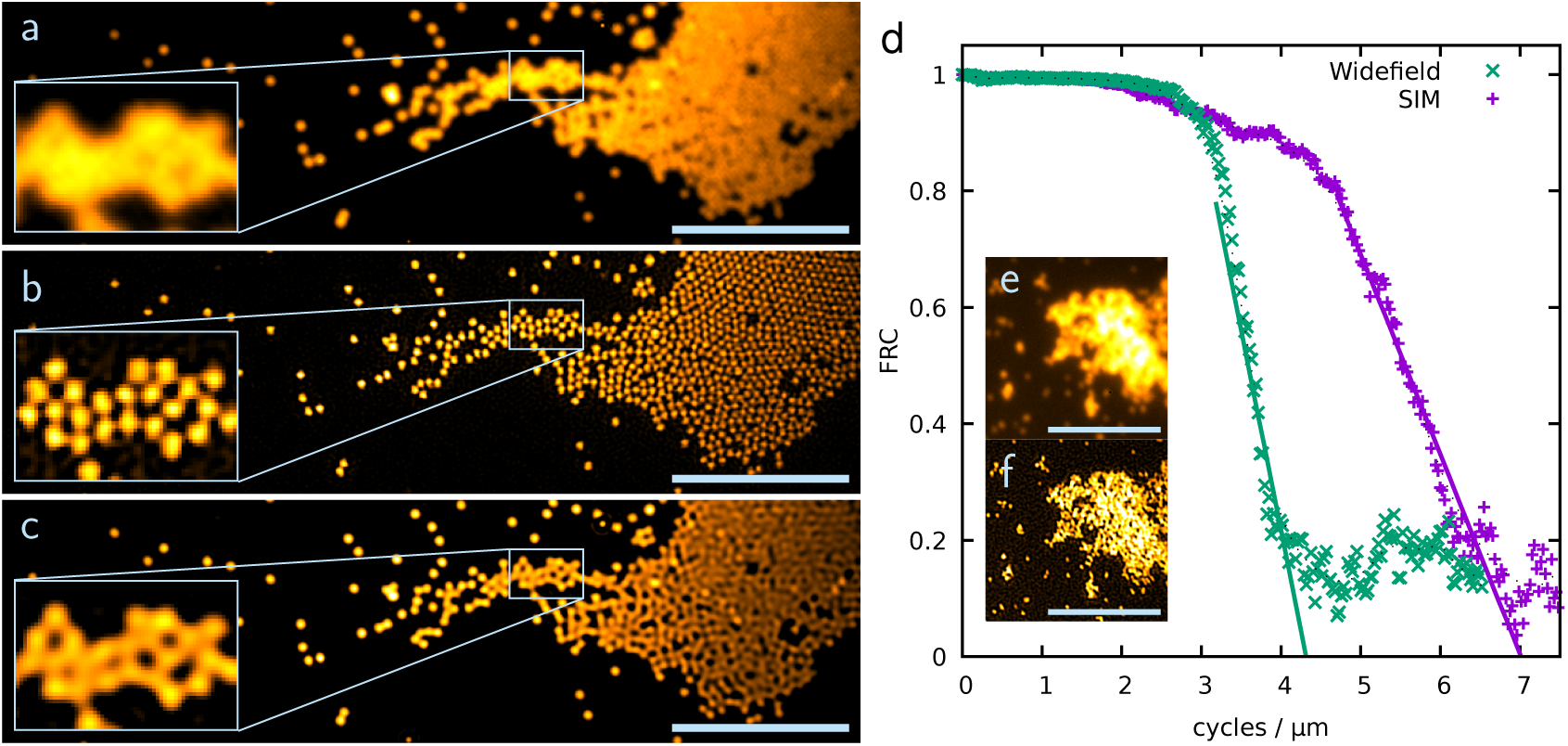
Fluorescence images of 200 nm TS beads (a-c) and FRC statistics (d). (a) Summed up wide-field image of the recorded nine SIM frames and filtered wide field image (c) determined by Wiener filtering. (b) Reconstructed SIM image of the 200 nm TS beads by fairSIM [30]. The inset clearly shows that single beads can be distinguished through SIM (scale bar 5 *µ*m, inset 2.1 *µ*m x 1.4 *µ*m, exposure time per raw frame 20 ms). Quantification of the resolution enhancement via FRC analysis on 100 nm beads (see inset e-WF, f-SIM), which are approximate point sources better than 200 nm beads (scale bar 5 *µ*m, exposure time per raw frame 50 ms). Two successive SIM imaged were acquired, reconstructed and analyzed via[31, 32]. The resulting graph (d) for both wide-field and SIM can be interpreted as the available signal (y-axis) at any given structure size (x-axis). By fitting both WF and SIM, a shift of 1.62× is found, which corresponds well with the expected resolution enhancement of 1.75×.

The SIM reconstruction process also allows us to estimate the pattern modulation depth achieved by the instrument. Here, this estimate yields 0.75, 0.79 and 0.74 for the respective SIM angles, which are reasonable values for a well-aligned 2D SIM system. To further quantify the system performance, the lateral resolution gain was determined. To this end, we image 100 nm TS beads in order to apply a Fourier Ring Correlation (FRC) analysis[31, 35]. In brief, FRC takes two measurement of the same structure, but with independent noise contribution, as input. It then computes the correlation of those measurements at different spatial frequencies, assuming no directionality (hence averaging over ‘rings’). The resulting graph can be interpreted as the available signal level of the imaging method at any given spatial frequency, and thus yields a model-free indicator of resolution. By applying this method to the DMD-SIM microscope data, we find a steep, expected drop-off in both the WF and the SIM datasets towards their respective resolution limits. By fitting these drop-offs (fig. 8d), we find that the FRC curve for SIM images drops off at 1.62 × higher spatial frequency than the WF signal. The SIM pattern in use is set to approximately 280 nm line spacing, with should yield an approximately 1.75 × resolution improvement at perfect SNR. However, the high frequency component accessible through SIM is dampened by the not fully modulated SIM pattern. Thus, the estimated modulation depths, the experimentally determined resolution calculated from FRC, and the theoretically expected resolution calculated from the SIM pattern spacing and optical parameters are fully consistent.

## 4 SIM images of biological samples

To further demonstrate the full functionality of our DMD-SIM microscope, we also imaged biological samples. First, we tested the imaging quality on fixed cells stained with different fluorescent dyes. Subsequently, various intracellular organelles labeled by fluorescent proteins were used to demonstrate that different structures can be resolved in vitro with our SIM microscope. Lastly, to further ensure that this system proves useful in a biological laboratory setting, we checked its live-cell compatibility by imaging living cells stained with live-cell stains.

### 4.1. Resolving subdiffraction sized structures in fixed cells

As an initial sample, we used fixed U2OS cells embedded in glycerol and stained their actin cytoskeleton with Phalloidin Atto532. These filaments are perfect organelles to demonstrate the resolution enhancement (fig. 9). In the WF images closeby single filaments cannot be resolved and appear as a single line. When switching to SIM mode, however, it becomes quite apparent that these are composed of more than one filament. These data also show that our instrument achieves an even illumination distribution over a large field of view. Therefore, extended structures with subdiffraction details in more than one cell can potentially be observed and analyzed.

**Figure 9:**
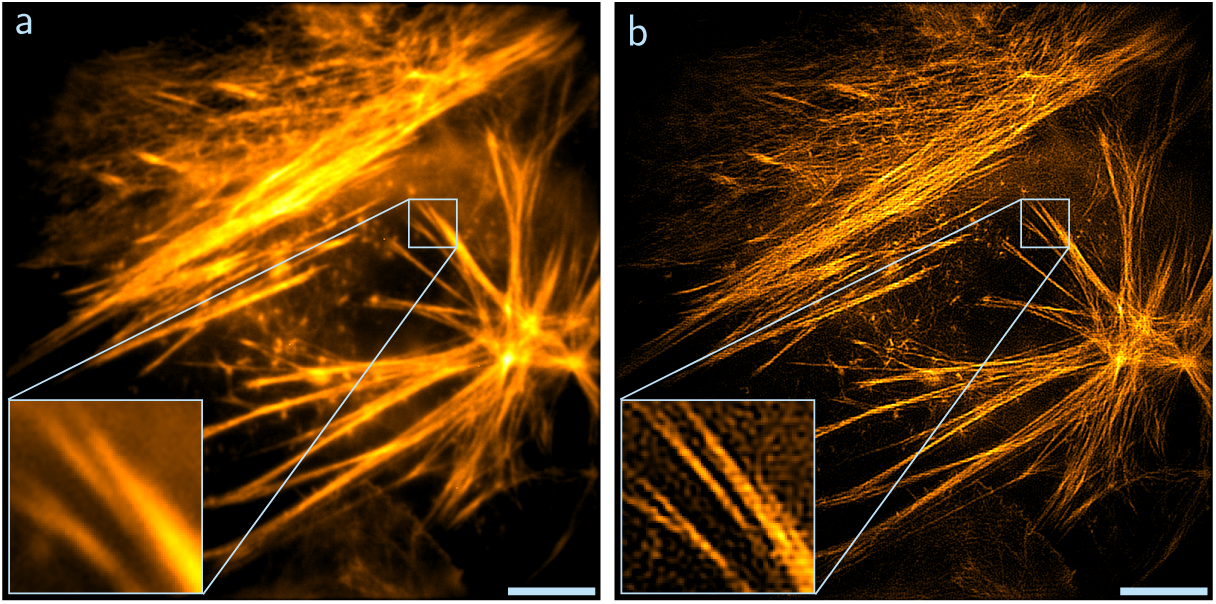
Fixed U2OS cell labeled with Phalloidin Atto532, WF (a) and SIM (b) image (exposure time per raw frame 20 ms). The size of the images is 36 *µ*m x 36 *µ*m which is also the possible field of view with the microscope. The actin filaments are now distinguishable in the SIM image (scale bar 5 *µ*m, inset 2.8 *µ*m × 2.8 *µ*m).

Next, we stained the actin cytoskeleton of fixed HEK293T cells with the red fluorescent protein mScarlet by gene transfection (fig. 10a and b). A direct comparison between fig. 9 and fig. 10 is not particularly suitable because the cell lines are different and mScarlet attaches to a different domain of the actin cytoskeleton than Phalloidin. Nonetheless, it is of interest to demonstrate SIM with mScarlet since fluorescent proteins are not as photostable as organic fluorescent dyes. Although a 532 nm laser is not the best choice for the excitation of mScarlett, our DMD-SIM microscope proves to have sufficient sensitivity to resolve the actin filaments which appear as single structures in the WF image.

**Figure 10:**
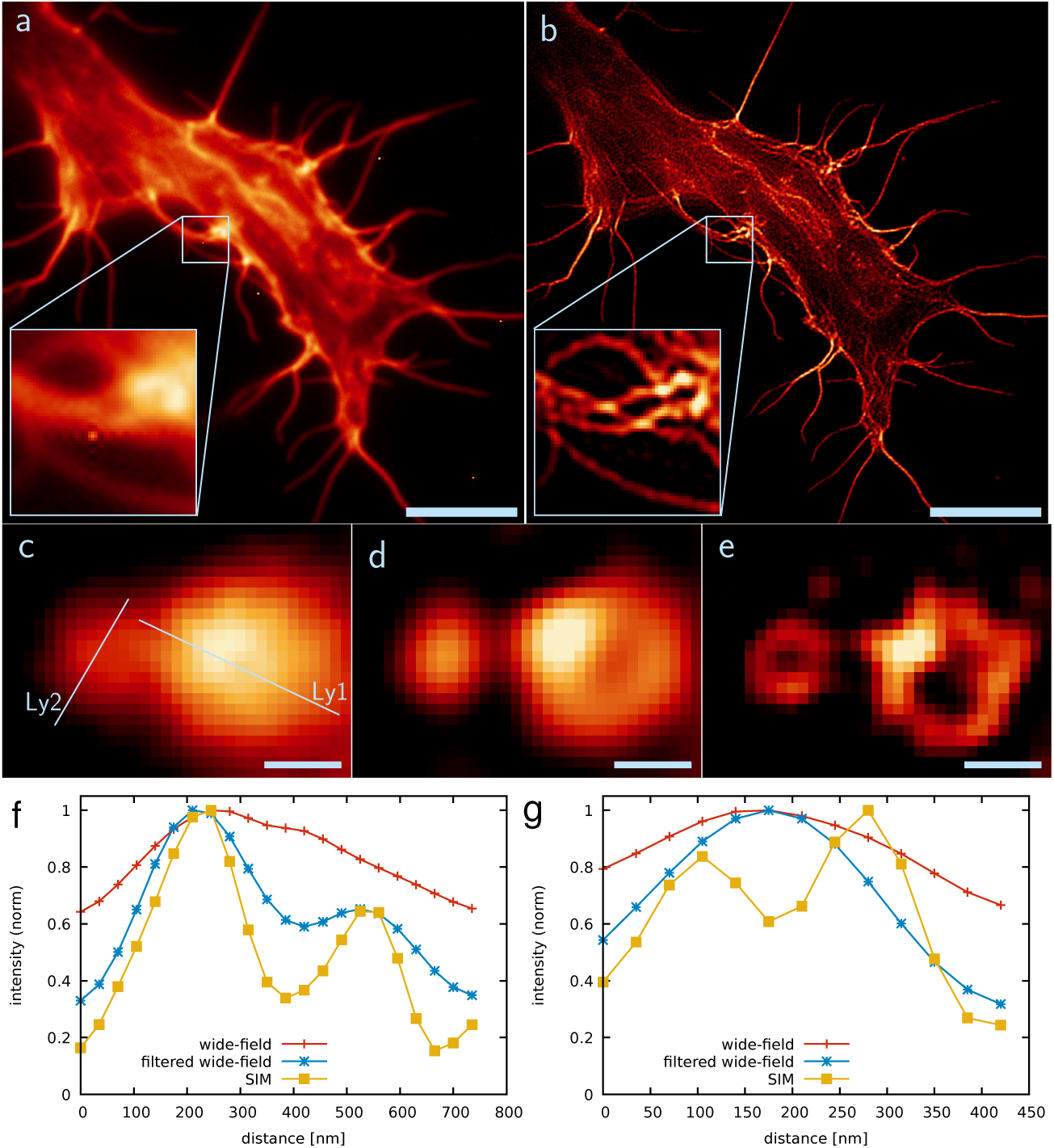
Fixed HEK293T cells transfected with the fluorescent protein mScarlet, to either label actin (a,b) or lysosomes (c-e). Actin filaments that are very close to each other cannot be distinguished in the WF image (a), but the SIM image reveals that more than one filament is present (b) (scale bar 5 *µ*m, inset 2.1 *µ*m x 2.1 *µ*m, exposure time per raw frame 50 ms). Lysosomes have different diameters and with the used plasmid, the outer membrane was stained. The membrane structure of bigger lysosomes (cross-section Ly1, plot e) can be revealed with conventional WF imaging (c) and additional filtering (d), but for smaller lysosomes (cross-section Ly2, plot f) SIM imaging is required (scale bar 250 nm, exposure time per raw frame 50 ms).

Furthermore, we labeled the outer membrane of lysosomes in HEK293T cells with mScarlet and subsequently fixed the cells. This organelle is of interest because it has a spherical shape and is present in a range of different diameters, from 100 nm to 1 *µ*m. Such spherical structures serve as a good quality check for SIM microscopes. If the resolution enhancement should be irregular in some directions, the organelle would appear elliptical rather than spherical. In addition, some lysosomes can be resolved via WF imaging but the smallest ones cannot. We demonstrate that our DMD-SIM microscope can resolve small spherical lysosomes (fig. 10c-g). In case the lysosome has a larger diameter than 250 nm, WF imaging is sufficient to resolve the outer membrane structure (fig. 10c and f). To underline this statement we apply fWF to increase the contrast, and this clearly shows that WF resolution is sufficient to resolve these organelles (fig. 10d and f). For structures with a diameter well below the diffraction limit, however, SIM is required and we can resolve lysosomes with a diameter of 180 nm which were not visible in WF (fig. 10e and g).

### 4.2. Live-cell imaging

In contrast to imaging fixed samples, live-cell imaging is more challenging. On the one hand, the refractive index of the medium surrounding the cells is different as they are now covered by water instead of being embedded in gycerol. On the other hand, the organelles are now mobile which might lead to motion-blur in the images. To circumvent these factors, we recorded image data at room temperature and utilized more processing for the image reconstruction. First, we subtracted background signal from the raw data in fairSIM in order to get rid of unwanted background contributions. To further enhance the contrast, fairSIM also allows us to apply Richardson-Lucy deconvolution to the input and output data [36]. Finally, by applying Hessian denoising to the reconstructed data it is possible to further smoothen the fluorescence signal and further enhance the contrast although some resolution improvement will be lost (parameters: *µ* = 100, *σ* = 0.8) [37].

For our first live-cell experiments we stained mitochondria in U2OS cells using the live cell stain MitoTrackerRed (fig. 11). This organelle has an inner structure, the so-called cristae, which is basically a folded membrane. With WF the cristae cannot be resolved because they are too close together, and therefore mitochondria are a popular organelle to demonstrate super-resolution microscopy. With our DMD-SIM microscope we recorded live-cell images of mitochondria. The summed up WF image reveals no inner structure (fig. 11a). However, at this point the real-time reconstruction with fairSIM-VIGOR was very convenient because it provided immediate feedback already during the data acquisition that confirmed that in SIM mode we were able to resolve the cristae. For further post-processing of the recorded data, we subtracted spurious background contributions since the dye also attaches in a small amount to other organelles. In addition, we applied Richardson-Lucy deconvolution to the raw input data to fairSIM, as well as the reconstructed output data, which reduces reconstruction artifacts. This results in a super-resolution image that resolves the inner structure of mitochondria quite well with a conclusive signal to noise ratio (fig. 11b). Nonetheless, by applying Hessian denoising to these data, the image quality can be further improved (fig. 11c).

**Figure 11:**
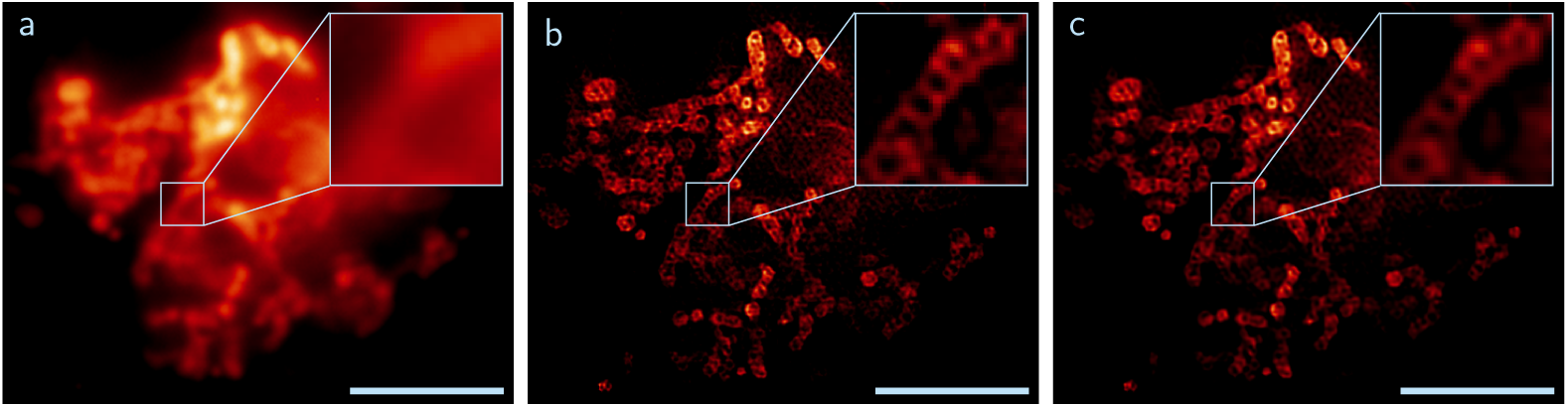
Live cell images of mitochondrial motility in U2OS cells. The organelle was stained with MitoTrackerRed and the data were recorded at room temperature. The insets show clearly that the inner structure, the cristae, cannot be resolved with WF imaging (a). For the SIM reconstruction, background signal from the raw data was subtracted and Richardson-Lucy deconvolution (10 iterations) was applied to the input and output images (b). Hessian denoising leads to further improvement of the image quality (c) (scale bar 5 *µ*m, inset 1.4 *µ*m × 1.4 *µ*m, exposure time per raw frame 100 ms).

To further demonstrate the live-cell imaging compatibility of our DMD-SIM microscope we recorded the movement of the endoplasmic reticulum (ER) network in U2OS cells (fig. 12). Here, the organelle was stained with ER-TrackerRed and we set a delay time of 250 ms between the SIM frames to keep the stress level for the cells to a minimum and to avoid photobleaching. Therefore, we were able to record live cell movies which last at least two minutes. Again, the real-time image reconstruction was very convenient for navigating the sample and finding sample areas of high motility. With SIM the dense ER network can be resolved and the movement of single filaments can easily be resolved (fig. 12a). Further image processing with Hessian denoising is very convenient in this case, because it smoothens the filaments and makes it easier to follow their movement (fig. 12b-d). In fig. 12c it is visible that one filament elongates along the x-axis. In addition, we can observe the detachment of a filament at a knot within the network and, direct following, its attachment to another point, which is a typical behavior for ER (fig. 12d, first row) [38]. Directly at this point the fiber then elongates over a long distance (fig. 12d, second row).

**Figure 12:**
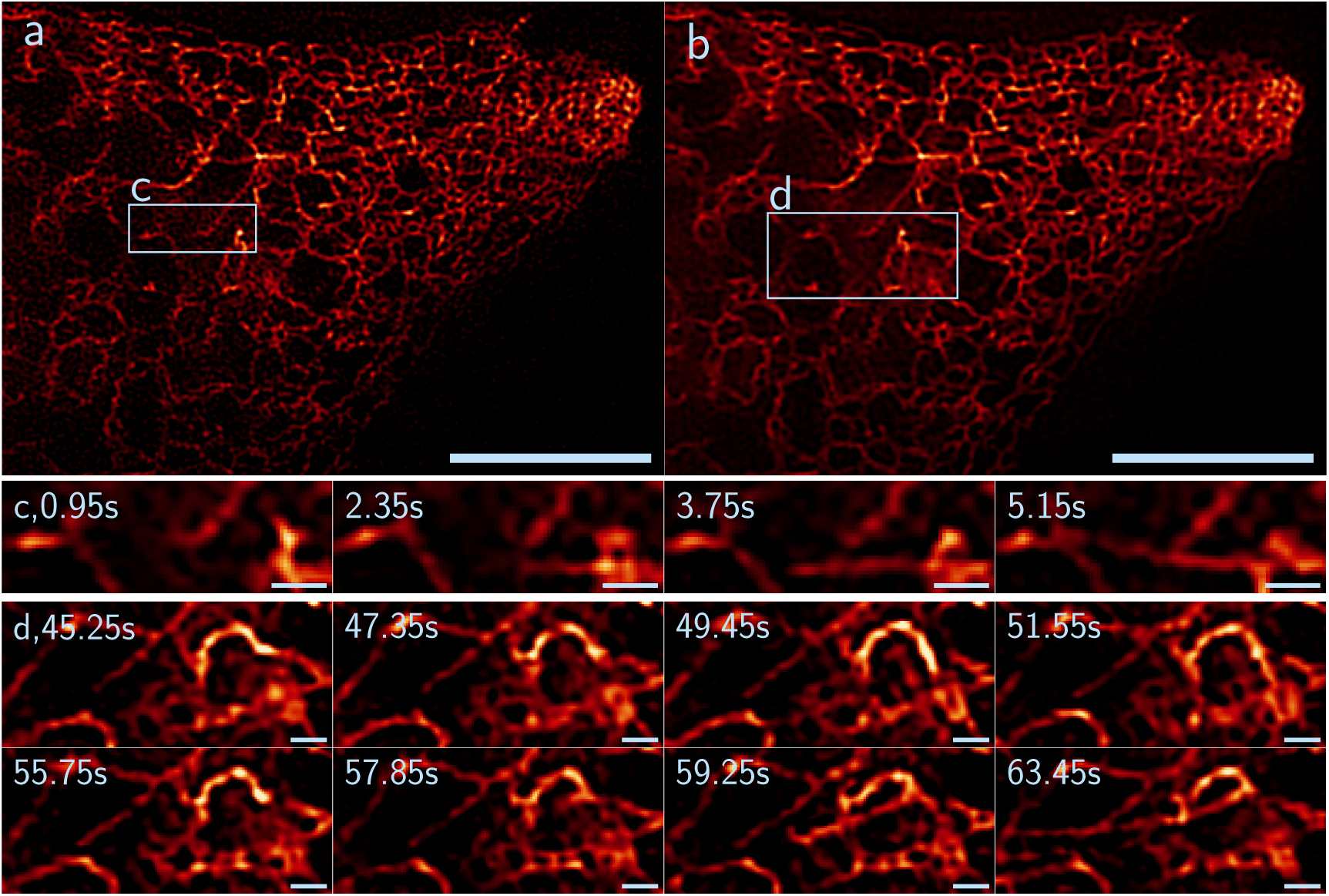
Endoplasmic reticulum (ER) network in a living U2OS cell. The organelle was stained with ER-Tracker Red and the data was recorded with a delay of 250n ms between each SIM sequence at room temperature. (a) SIM image at time point 0 s reveals many filaments. (b) Hessian denoising of the SIM frame reduces the background signal and smoothes the filaments (scale bar 5 *µ*m, exposure time per raw frame 50 ms). The time sequence (c) is at inset 1 and shows the elongation of a fiber (scale bar 500 nm). In addition, a detachment can be observed with a subsequent attachment to another point and a further elongation (d, inset 2, scale bar 500 nm). Both insets display the Hessian denoised data.

## 5. Discussion and conclusion

Our work provides two significant contributions that further facilitate the overall use of digital micromirror devices for coherent illumination, which is required to utilize structured illumination microscopy to its fullest potential. Our simulations provided specific information on how the blazed grating effect caused by the DMD substructure affects the SIM illumination pattern, how to identify ideal alignment angles for the device, as well as how manufacturing tolerances in the device affect this alignment. Our design of a DMD-SIM system then utilized these findings, and it enabled us to employ a DMD as the light modulator resulting in a very compact and cost-effective SIM system. We showcased its capabilities by providing calibration measurements, as well as conducting fixed and live-cell imaging at sub-second acquisition times.

We view this work as a contribution to the growing efforts of democratizing super-resolution microscopy. Digital micromirror devices are both cost-effective and readily available in many variations, which makes them an attractive alternative to the specialized SLM devices typically employed for SIM, which require more difficult to implement timing schemes [10, 11]. By providing a thorough characterization of the blazed grating effect in the context of SIM, it should now be easy to employ DMDs in different and more complex variations of novel SIM systems.

Following the spirit of open science, we provide all source code and raw data for the results presented in this manuscript to the scientific community. All code is openly accessible under GPLv2 (or later) license, and can be found in online repositories under github.com/fairSIM and github.com/biophotonics-bielefeld.

## Author contributions

A.S. built the DMD-SIM microscope, performed the blazed grating effect and SIM measurements, processed the SIM reconstructions, prepared samples, and wrote the manuscript. M.L. wrote the DMD simulation software, set up electronics and control software of the microscope, and wrote parts of the manuscript. H.S. helped to implemented and crosschecked math and simulation of the blazed grating effect. W.H. prepared the fixed and live-cell samples. T.H. supervised the project, and helped write the manuscript. M.M. conceived of and supervised the project, performed FRC analysis, and wrote the manuscript. All authored took part in editing and approve of the manuscript.

## Acknowledgments

We would like to thank Frederik Lange, Mick Phillips, Ian Dobbie and Rainer Heintzmann for fruitful discussions of the blazed grating effect applied to SIM illumination. We also like to thank Andreas Markwirth for his help in setting up the control electronics and Jochen Linnenbrügger for constructing the DMD holder. We acknowledge funding by the Deutsche Forschungsgemeinschaft (DFG, German Science Foundation) - project number 415832635. This project has also received funding from the European Union’s Horizon 2020 research and innovation programme under the Marie Skłodowska-Curie grant agreements No. 752080.

## A. Detailed description of the SIM microscope

### A.1. Illumination path and data acquisition

The 532 nm diode laser is expanded (f(L1) = 30 mm, and f(L2) = 150 mm) and then coupled into a high-power fiber via a 10x objective. The laser power level cannot be controlled electronically, and therefore two neutral density filter (10 % and/or 50 %) are utilized if the laser power needs to be adjusted. With a customized collimator (f(L3) = 20 mm) the laser beam is paralleled and directed to the DMD with the blazed angle as the incident angle. The DMD itself is also mounted on a customized holder which a tilt of 45°. The holder can shift the DMD in x- and y-direction and adjust the tilt along the xz- and yz-plane. The L4 lens (f = 200 mm) collects the diffracted light and a combination of polarization filter and *λ*/4 plate converts the elliptic polarized light to circular polarized light. In the Fourier plane of L4, only the first side orders of the main diffracted orders pass the Fourier filter. Directly afterwards these laser beams are linearly polarized by a “pizza”-polarizer, so that the side orders of the same SIM angle have the same linear polarization. A L5 lens (f = 100 mm) collects the beams and the first dichroic mirror reflects them to L6 (f = 150 mm) which is focusing the light in the back-focal plane of the objective. A dichroic mirror from the same batch as the first one, reflects the laser beam to the immersion oil objective which focuses the SIM beams in the sample. For the data acquisition the Zeiss 1.518 oil is used and a xyz-block is used for the focusing and maneuvering of the sample. The objective collects the fluorescence signal which then passes the second dichroic mirror and is reflected in direction to the camera. A detection filter is implemented to block unwanted laser light. Either an orange detection filter is used or a red detection filter, suitable to the used dye. The lens L7 (f = 250 mm) is used to focus the fluorescence emission on the camera. System synchronization, data acquisition and real-time SIM image processing is achieved through the fairSIM-VIGOR engine[11] and the MicroManager software package[39, 40].

### A.2. Component list

The main used items for our DMD-SIM microscope can be found in table 1. All the other commercial optomechanics are basic components from Thorlabs like the cage system. Here, we utilized for instance construction rods, mounting brackets and posts.

**Table 1:**
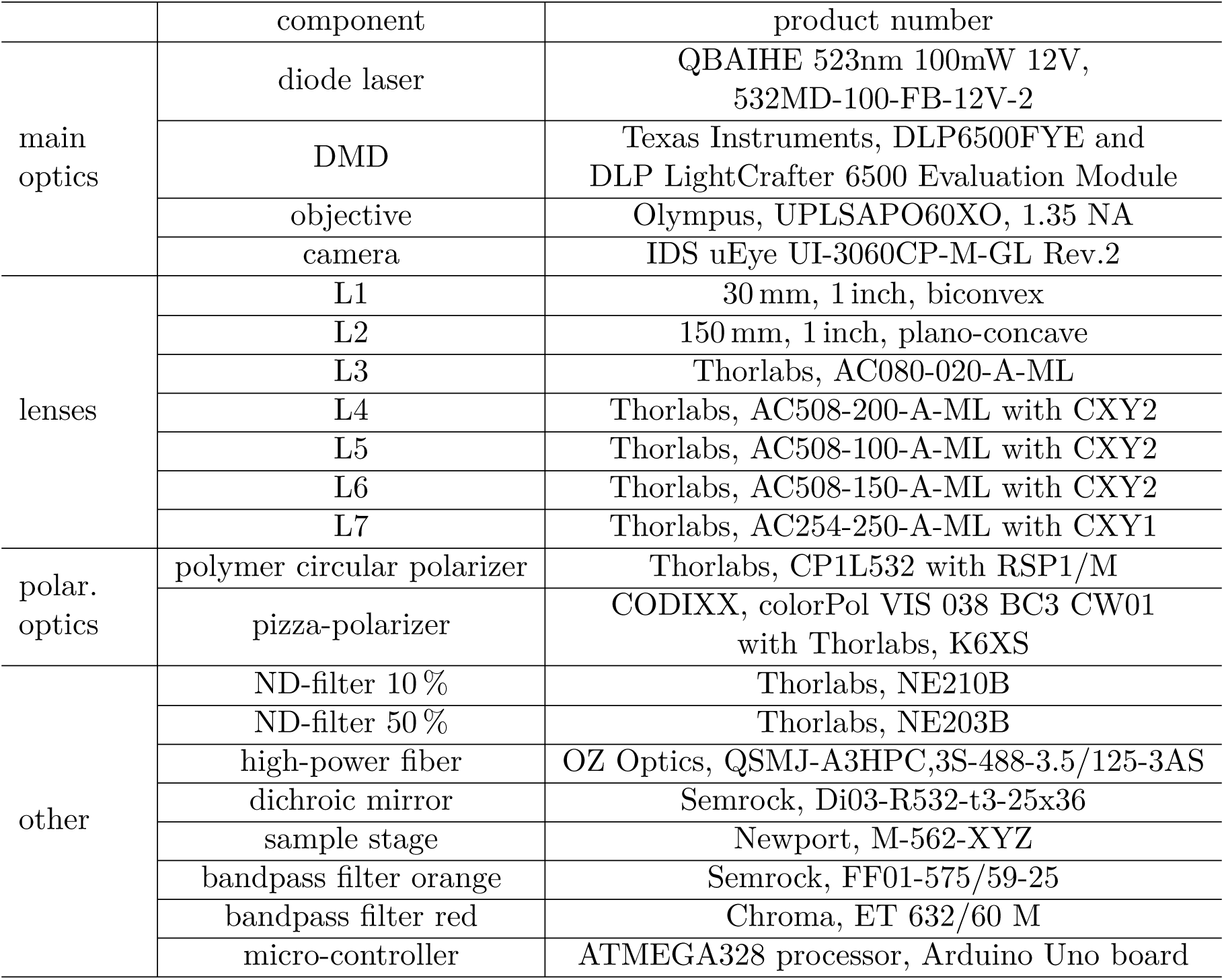
List of the most important components of the DMD-SIM setup. The position of the optics and description of the lenses can be seen in fig. 7a.

## B. Blazed grating angle determination software

A software package to simulate the blazed grating effect and determine the intensity distributions depending on the angle of incidence, light wavelength and SIM pattern has been created. The software is written in Java to work as a plugin to scientific image processing package ImageJ[41]/FIJI[42]. and provides GPU acceleration through the jCuda framework[43]. The source code is available under an open-source (GPLv3) licence in an online repository: github.com/biophotonics-bielefeld/coherent-dmd-sim-simulator.

## C. Sample preparation

### C.1. Preparation of TetraSpeck bead slides

For the bead slides we purchased TetraSpeck Microspheres from Thermo Fischer in the sizes 200 nm (#T-7280) and 100 nm (#T-7219). Both bead slides have been prepared the same way and will be addressed as TS. The TS stock solution was diluted with double distilled water down to a 1:10 concentration. A small droplet of approximately *µ*l was placed on a ∅12 mm coverslip and spread all over it with the pipette tip. After an incubation time of 1 h at room temperature in the dark, the droplet was dried. Now, *µ*l of glycerol was added on the coverslip and a cover glass was placed on top without getting bubbles in the glycerol. Lastly, the sample has been sealed with nail polish at the edge of the coverslip on the cover glass.

### C.2. Biological samples

All the cells were grown on laminin-1 (Sigma-Aldrich, #L2020) coated #1.5 high precision glass. A CO_2_ independent medium (Gibco, #18045088) was used for live microscopy experiments.

#### Fixed cells

HEK293T cells were transfected for transcient expression of LifeAct-mScarlet and vLamp1-mScarlet fusions with Lipofectamine 3000 (ThermoFisher, #L3000015) according to the manufacturer’s protocol. pLifeAct-mScarlet-N1 and Lamp1-mScarlet-I were a gift from Dorus Gadella (Addgene plasmid # 85054 ; http://n2t.net/addgene:85054 ; RRID:Addgene_85054 and Addgene plasmid # 98827 ; http://n2t.net/addgene:98827 ; RRID:Addgene_98827) [44, 45]. The cells were fixed 24 h post-transfection with 4 % paraformaldehyde at room temperature for 10 min.

#### Living cells

U2OS cells were stained for mitochondria or endoplasmic reticulum with respectivily Mi-toTracker Red (ThermoFisher, #M22425) or ERTracker Red (ThermoFisher, #E34250) according to the manufacturer’s protocols.

